# Why Does Synergistic Activation of WASP, but Not N-WASP, by Cdc42 and PIP_2_ Require Cdc42 Prenylation?

**DOI:** 10.1101/2022.11.24.517863

**Authors:** Souvik Dey, Huan-Xiang Zhou

**Author notes:** Correspondence should be addressed to H.X.Z.

## Abstract

Human WASP and N-WASP are homologous proteins that require the binding of multiple regulators, including the acidic lipid PIP_2_ and the small GTPase Cdc42, to relieve autoinhibition before they can stimulate the initiation of actin polymerization. Autoinhibition involves intramolecular binding of the C-terminal acidic and central motifs to an upstream basic region and GTPase binding domain. Little is known about how a single intrinsically disordered protein, WASP or N-WASP, binds multiple regulators to achieve full activation. Here we used molecular dynamics simulations to characterize the binding of WASP and N-WASP with PIP_2_ and Cdc42. In the absence of Cdc42, both WASP and N-WASP strongly associate with PIP_2_-containing membranes, through their basic region and also possibly through a tail portion of the N-terminal WH1 domain. The basic region also participates in Cdc42 binding, especially for WASP; consequently Cdc42 binding significantly compromises the ability of the basic region in WASP, but not N-WASP, to bind PIP_2_. PIP_2_ binding to the WASP basic region is restored only when Cdc42 is prenylated at the C-terminus and tethered to the membrane. This distinction in the activation of WASP and N-WASP may contribute to their different functional roles.

## Introduction

Human Wiskott-Aldrich syndrome protein (WASP) and neural WASP (N-WASP) are homologous proteins that contain intrinsically disordered regions throughout their sequences [1]. Typically of intrinsically disordered proteins (IDPs), WASP and N-WASP engage in numerous intramolecular and intermolecular interactions. Intramolecular interactions result in autoinhibition [2-5] whereas intermolecular interactions result in activation and subsequent engagement in cellular activities. A major activity of WASP and N-WASP is to stimulate the initiation of branched actin polymerization [6]. WASP is expressed only in hematopoietic cells [7], whereas N-WASP, despite its prefix, is ubiquitously expressed [8]. Mutations in WASP are responsible for its namesake, an X-linked immunological disorder [7]. The fact that N-WASP in the hematopoietic cells of patients does not compensate for the defects caused by WASP mutations points to nonredundant roles of these two proteins. Indeed, different effects of WASP and N-WASP have been reported in filopodium formation [9], podosome-mediated extracellular matrix degradation [10], and T cell chemotaxis [11]. Phylogenetic analysis suggested a direct link between N-WASP and ancestral WASPs, whereas hematopoietic WASP branched off during the onset of vertebrates [12]. WASP and N-WASP are excellent models for studying how a single IDP binds multiple regulators to achieve full activation as well as for deciphering the differences between closely related IDPs in processes ranging from activation to cellular activities.

WASP and N-WASP share ∼50% sequence homology [8] and have the same domain organization with one exception [1, 13] (Figure 1A). The functional element resides toward the C-terminus, where the verprolin-central-acidic (VCA) region binds G-actin (via the V motif) and Arp2/3 (via the CA motifs) to initiate actin polymerization [2, 3, 14, 15]. The exception to the shared domain organization is an additional V motif in N-WASP, which increases the activity for stimulating actin polymerization [16]. Upstream of the functional element are regulatory elements. At the N-terminus is a WASP homology 1 (WH1; also known as EVH1) domain that binds WASP-interacting proteins [17]. Following a linker of ∼65 (or ∼35 in N-WASP) residues, a basic region (BR) is present to bind the acidic lipid phosphatidylinositol 4,5-bisphosphate (PIP_2_) [3, 5, 18-20]. Higgs and Pollard [18] suggested that the WH1 domain of WASP may also participate in PIP_2_ binding. Immediate after the BR is a GTPase-binding domain (GBD), which binds Cdc42 and other small GTPases [9, 21]. In WASP, several Lys residues in the basic region, including K_230_KK_232_, are involved in electrostatic interactions with Cdc42 [22] and contribute to their binding rate [23-25]. Lastly, a proline-rich region (PRR) connects the GBD and the VCA region, and binds SH3-containing proteins such as Nck [26].

**Figure 1.**
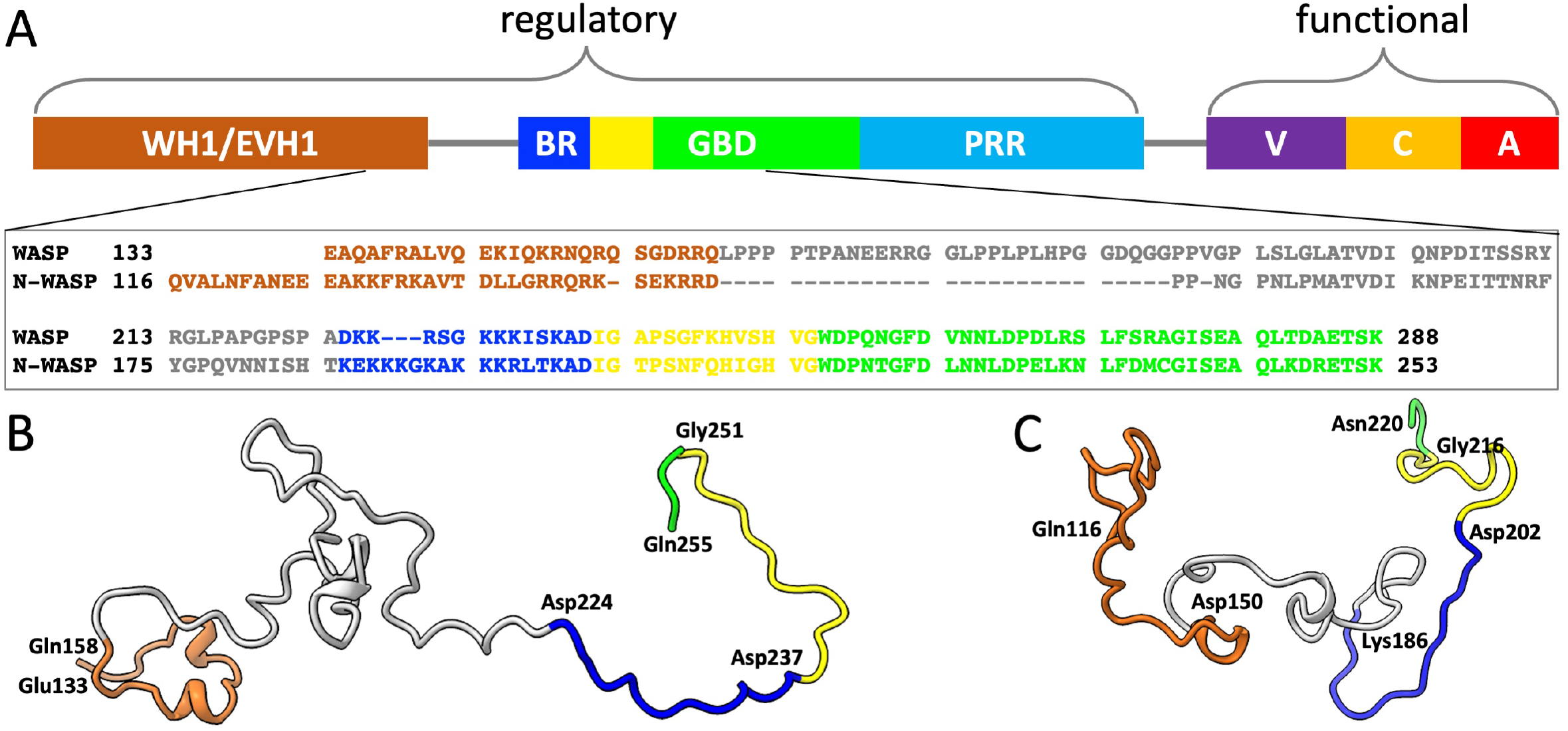
Sequences and conformations of WASP and N-WASP. (A) Domain organization of WASP and sequence alignment of WASP and N-WASP for residues implicated in PIP_2_ and Cdc42 binding. The domains and disordered regions are indicated by different colors. The initial portion of the GBD is a CRIB (Cdc42/Rac interactive binding) motif, colored yellow; the rest of the GBD is colored green. The domain organization is shared by N-WASP, with two exceptions: the linker between WH1 and BR is shorter by ∼30 residues, and an extra V motif is inserted between PRR and VCA. The sequences are color coordinated with the domain organization. Representative conformations of (B) WASP and (C) N-WASP while bound to PIP_2_-containing membranes. Colors of different regions are as in (A); the bordering residues between regions are labeled.

Under resting conditions, WASP and N-WASP are autoinhibited by intramolecular interactions, between the GBD and the C motif [3-5, 9] and between the BR and the A motif [1, 25, 27]. Binding to PIP_2_ and Cdc42 relieve N-WASP of autoinhibition, and the two regulators have synergistic effects on activation [3]. Endogenous Cdc42 is prenylated at the C-terminus and then tethered to membranes [28], but a follow-up study reported no difference between prenylated and soluble Cdc42 in their synergistic effects with PIP_2_ on N-WASP activation [19]. Prehoda et al. [5] also observed synergistic activation of N-WASP by PIP_2_ and soluble Cdc42, though a follow-up study found higher activity when Cdc42 was prenylated and colocalized with PIP_2_ [20]. In contrast, synergistic activation of WASP by PIP_2_ and Cdc42 required Cdc42 prenylation [18]. Indeed, instead of doubling the effect of PIP_2_ when prenylated Cdc42 was added, a 30% decrease was observed when soluble Cdc42 was added. Both the synergistic effect of soluble Cdc42 on N-WASP activation and the negative effect of soluble Cdc42 on WASP activation were reproduced by Tomasevic et al. [29], but the reason for this contrast has remained a mystery.

The structures of the WASP GBD bound to the C motif (modeling the autoinhibited state) and to Cdc42 (modeling the activated state) have been determined by NMR spectroscopy [4, 22]. In the autoinhibited state, the C motif folds into an α-helix and docks to a three-helix platform formed by the GBD (Figure S1A). In the activated state, the platform falls apart as one of these helices, along with upstream segments, binds to Cdc42 (Figure S1B). Very little structural information is available for the binding of WASP or N-WASP with the other activator, i.e., PIP_2_. Binding studies [20] have shown that the N-WASP BR binds multiple PIP_2_ molecules and it is the number of basic residues, not the precise sequence, that determines the PIP_2_ binding affinity, typical of IDP binding to acidic membranes [30]. Structural characterization of IDP-membrane binding, due to its dynamic nature, presents challenges for experimental techniques, but this problem can now be addressed by molecular dynamics (MD) simulations [31-34].

Here we report MD simulation results on the binding of WASP and N-WASP with PIP_2_ and soluble and prenylated Cdc42. The simulations provided an opportunity to visualize how WASP and N-WASP simultaneously bind with two regulators, PIP_2_ and Cdc42, thereby fully releasing the VCA region for stimulating actin polymerization. For N-WASP, simultaneous binding can occur whether Cdc42 is soluble or prenylated.

However, WASP can do so only when Cdc42 is prenylated. This result explains the previous puzzling observation that synergistic activation of WASP, but not N-WASP requires Cdc42 prenylation. The origin for this contrast lies in the differences between the BRs of WASP and N-WASP, in particular regarding the number of basic residues and their spacing from the GBD.

## Results

We simulated WASP and N-WASP fragments binding to PIP_2_-containing membranes in the absence of Cdc42 and in the presence of soluble and prenylated Cdc42. The upper leaflet (facing the proteins) contained 10% PIP_2_ and 40% POPS; the remaining 50% was neutral lipids. The simulations for WASP or N-WASP without Cdc42 were run in 16 replicates, each lasting 1360 ns. The simulations with either soluble or prenylated Cdc42 were run in four replicates, each lasting 1000 ns for WASP and 700 ns for N-WASP. We quantify membrane binding by calculating the membrane contact probabilities of individual residues, defined as the fraction of MD frames in which a residue forms at least one contact with lipids. A contact is considered formed when two heavy atoms come within 3.5 Å.

### The N-WASP BR shows stronger membrane association than the WASP counterpart

For the simulations without Cdc42, the WASP and N-WASP fragments contained residues Glu133 to Gln255 and Gln116 to Asn220, respectively. They start from the tail portion of the WH1 domain and end just beyond the CRIB motif of the GBD. A snapshot for each fragment from the simulations is shown in Figure 1B, C. The WASP and N-WASP BRs consists of 14 and 17 residues (Asp224 to Asp237 and Lys186 to Asp202; Figure 1A), with 7 and 10 of them, respectively, being basic. As shown in Figure 2A, B, the BRs have high membrane contact probabilities, with basic residues making dominant contributions. All the basic residues have >10% membrane contact probabilities; the only other residue crossing this threshold is Ser228 in the WASP BR.

**Figure 2.**
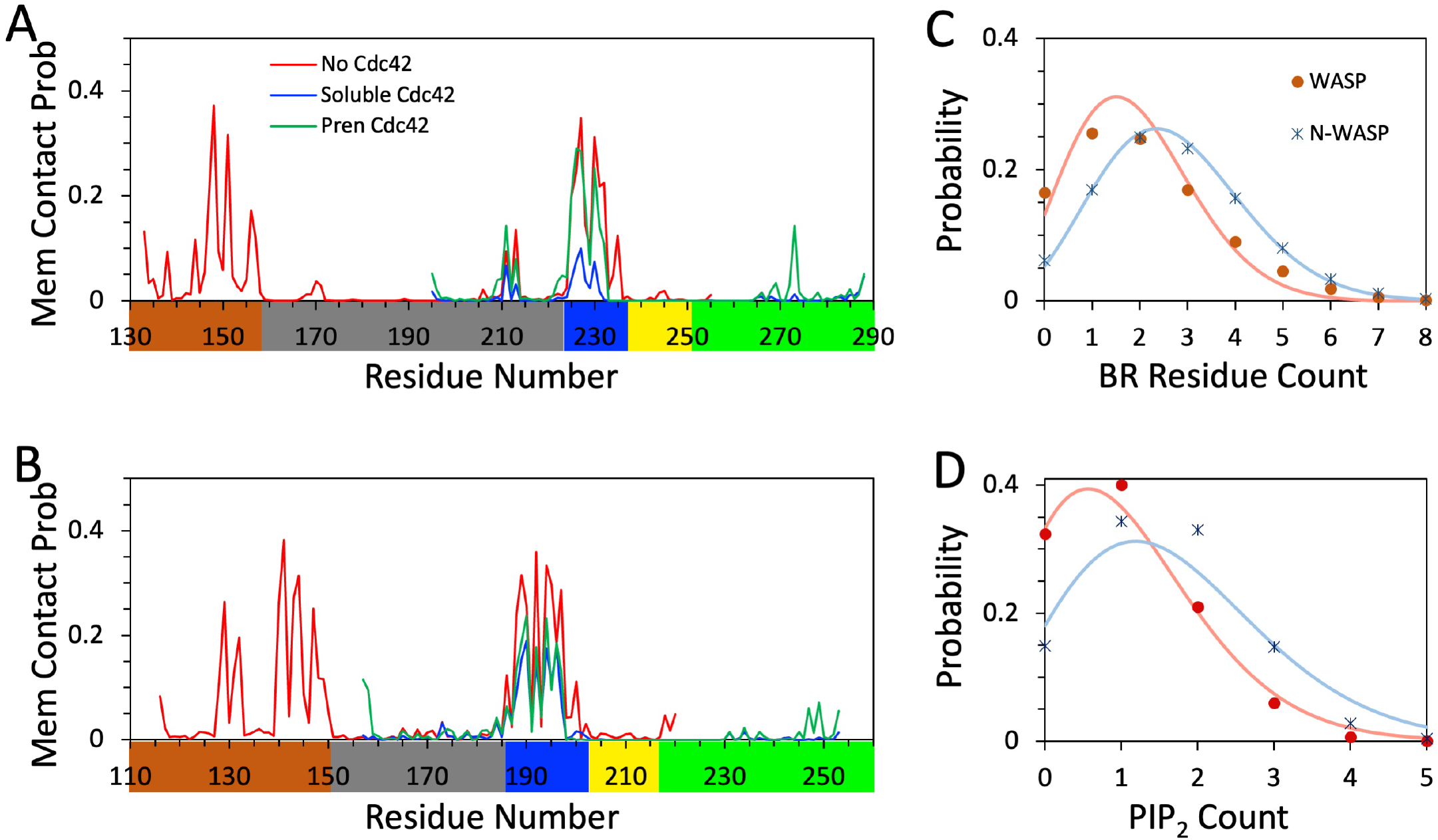
Membrane binding properties of WASP and N-WASP. Membrane contact probabilities of (A) WASP and (B) N-WASP residues. Traces in red, blue, and green display data from simulations without Cdc42, with soluble Cdc42, and with prenylated Cdc42, respectively. (C) Probabilities for having *n* BR residues simultaneously bound to membranes in simulations without Cdc42; *n* is BR residue count. (D) Probabilities for having *m* PIP_2_ lipids simultaneously bound to the BR in simulations without Cdc42; *m* is PIP_2_ count. In (C, D), data from the simulations of WASP and N-WASP fragments are shown as red and blue symbols, respectively; fits to a binomial distribution in (C) and Poisson distribution in (D) are displayed as curves in colors matching the corresponding symbols.

Because of its larger number of basic residues, the N-WASP BR shows stronger membrane association than the WASP counterpart. This difference can be measured in two ways. First, the fraction of MD frames where the BR is membrane-associated is higher for N-WASP; it is 92%, compared to 83% for WASP. Second, in a given frame, more residues in the N-WASP BR are simultaneously membrane-bound than in the WASP BR. Figure 2C displays the probabilities of having exactly *n* BR residues bound to membranes at a given time, with *n* = 0, 1, 2, … Relative to WASP, the probability distribution of the N-WASP shifts toward higher *n*. Both of the probability distributions fit moderately well to a binomial distribution,

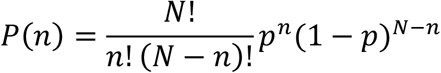

which assumes that the *N* BR residues (14 and 17 for WASP and N-WASP, respectively) each independently bind to the membrane with probability *p*. The predicted mean *n*, 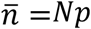, represents the average number of simultaneously bound BR residues. The fits yield *p* = 0.135 for WASP and 0.159 for N-WASP. The resulting mean *n* is 1.9 for WASP and 2.7 for N-WASP. The higher value for N-WASP is due to both a longer BR and a higher membrane binding probability for each BR residue.

Most of the lipids interacting with the basic residues are PIP_2_; POPS lipids also participate in the interactions to some extent. Figure 2D displays the probabilities of having exactly *m* PIP_2_ lipids simultaneously bound to the BR. The probability distribution in *m* also shifts toward higher *m* values for N-WASP – just as more N-WASP BR residues participate in membrane interactions, we see more PIP_2_ lipids simultaneously bind to this BR. The probability distributions in *m* are close to a Poisson distribution,

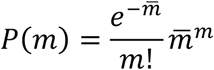

which is the limit of the binomial distribution when *N* → ∞ while *Np* remaining at a constant value 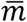. The closeness to a Poisson distribution means that PIP_2_ molecules also largely work independently when binding to the BRs. This outcome is perhaps befitting to the relatively low level of PIP_2_ (10%) in the membranes. The mean 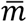 is 1.6 for N-WASP, compared to 1.0 for WASP.

Figure 3A displays a representative snapshot of the WASP fragment bound to the membrane. Several basic sidechains in the BR project downward and reach for lipid headgroups. The two BR sidechains with the highest membrane contact probabilities are Arg227 and Lys230 (Figure 2A). In the representative snapshot (Figure 3A, zoomed view), Arg227 forms a salt bridge with a PS headgroup while Lys230 hydrogen bonds with a PIP_2_. Two other sidechains, Lys232 and Lys235, are just outside the cutoff distance of 3.5 Å from lipids, and ready to form alternative or additional contacts with lipids.

**Figure 3.**
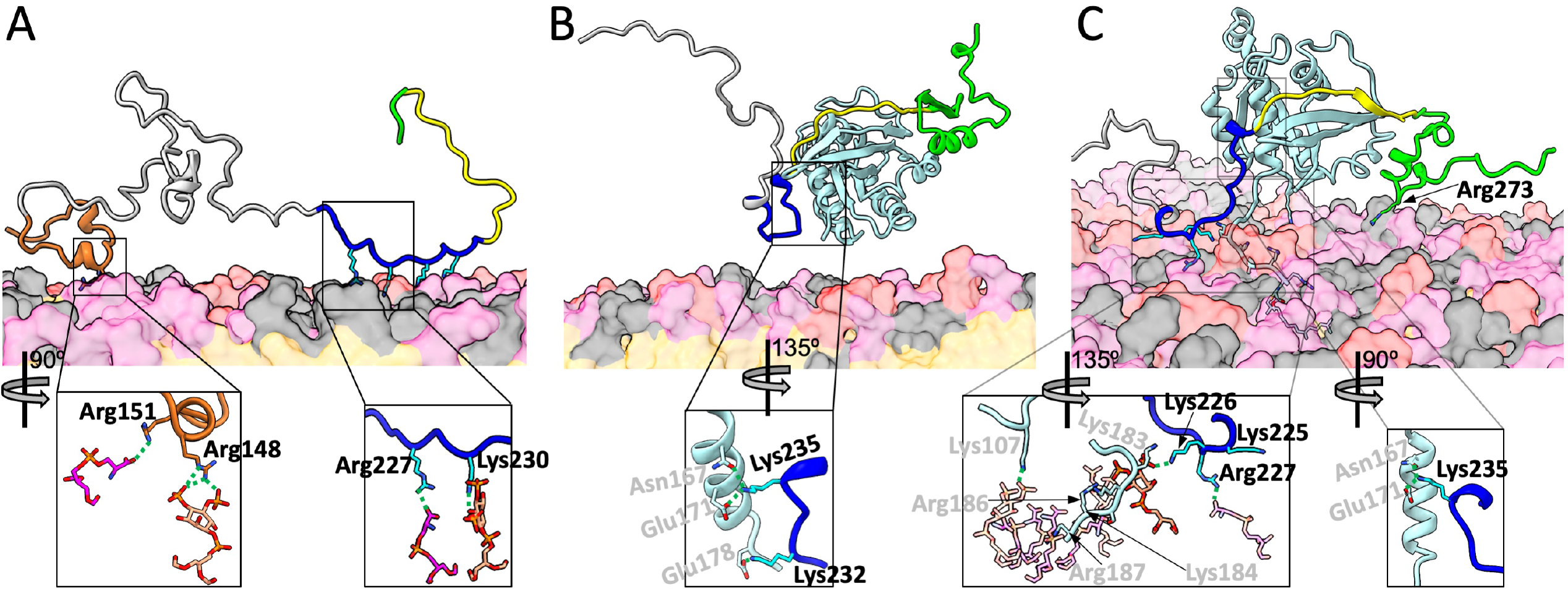
Representative snapshots from MD simulations of membrane binding of WASP. (A) Without Cdc42. (B) With soluble Cdc42. (C) With prenylated Cdc42. WASP residues are colored according to the scheme in Figure 1; Cdc42 is in light cyan. Lipids are shown as surface, with headgroups of PIP_2_, POPS, and neutral lipid (POPC, POPE, and cholesterol) in pink, magenta, and grey respectively; lipid tails are in orange. WASP sidechains that form or are about to form membrane contacts are shown as sticks. In the zoomed images, the lipid partners are also shown as sticks, as are BR-Cdc42 interactions. Residue labels are in black for WASP and gray for Cdc42, respectively. Carbon atoms have the same colors as the backbone representations, except that those in the BR are in cyan; O, N, and P atoms are in red, blue, and orange, respectively. In (C), the zoomed left image shows all but one lipid with dimmed intensity; the exception is a PIP_2_ that interacts with both WASP Lys226 and Cdc42 Lys183.

A representative snapshot of the N-WASP fragment is shown in Figure 4A. In this BR, the three sidechains with the highest membrane contact probabilities are Lys189, Lys192, and Lys194 (Figure 2B). In the representative snapshot (Figure 4A, zoomed view), Lys189 forms a salt bridge with a PIP_2_, Lys192 hydrogen bonds with a PIP_2_, and Lys194 forms a salt bridge with a PS. In this case, Lys195 is waiting to get closer to lipids.

**Figure 4.**
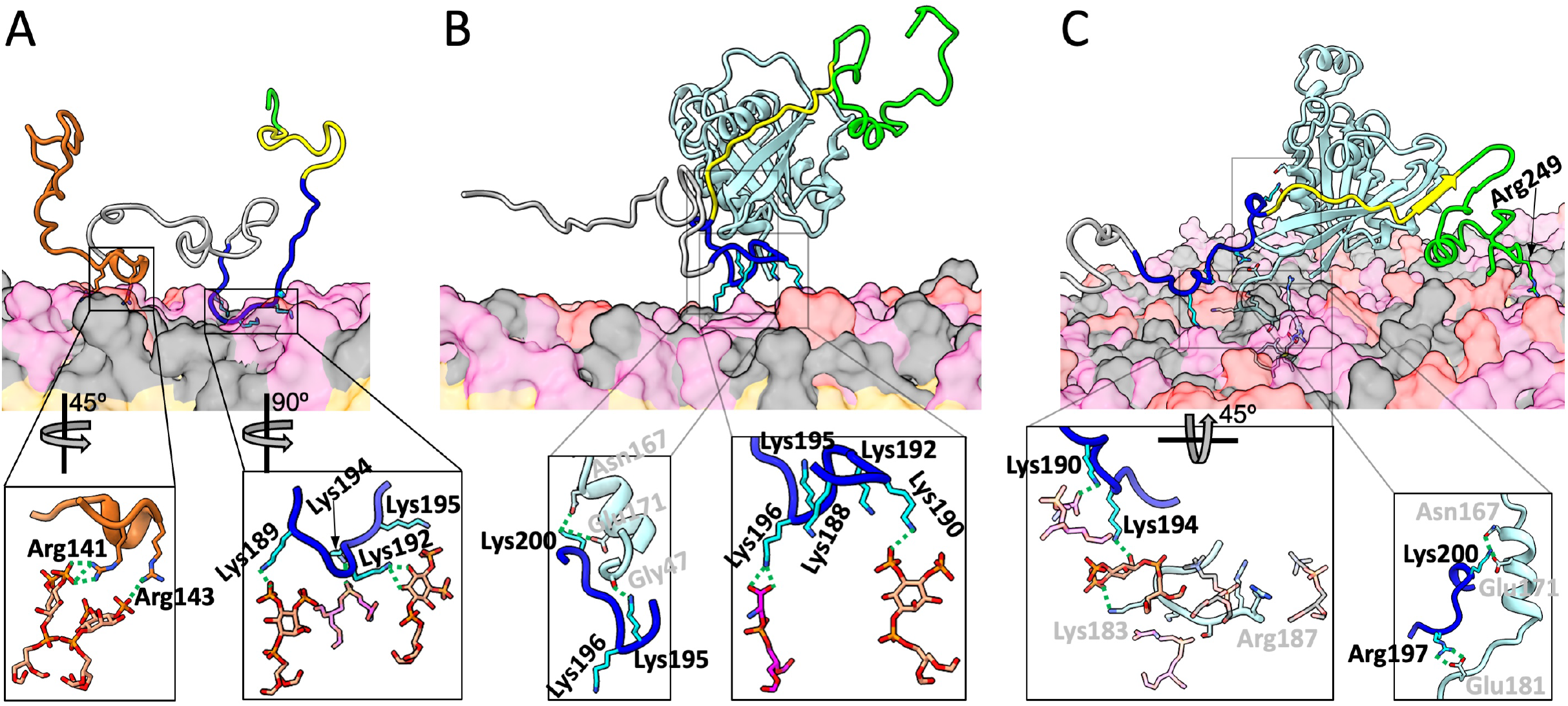
Representative snapshots from MD simulations of membrane binding of N-WASP. (A) Without Cdc42. (B) With soluble Cdc42. (C) With prenylated Cdc42. The representations and colors are the same as described in Figure 3, except that in (C), two salt bridges and a hydrogen bond between N-WASP and Cdc42 are shown in both the normal and zoomed views.

### A tail portion of WH1 may assist with membrane association

We included a portion of the WH1 domain in the WASP and N-WASP fragments for membrane association simulations in the absence of Cdc42, partly because Higgs and Pollard [18] suggested that the WASP WH1 may also participate in PIP_2_ binding. There was also an additional reason. We developed a sequence-based method called ReSMAP for predicting residue-specific membrane association propensities of IDPs, trained on MD simulation data including those from an initial portion of the WASP and N-WASP simulations [30]. We used an initial version of ReSMAP to select the WASP and N-WASP fragments for membrane association simulations. It already predicted significant membrane association propensities for the tail portions of the WASP and N-WASP WH1, due to a large number of basic residues there (7 and 11, respectively; Figure 1A). Hence the tail portions were included in the WASP and N-WASP fragments for membrane association simulations.

As shown in Figure 2A,B, the WH1 tail portions indeed show high membrane contact probabilities in the MD simulations. All but one basic residue (WASP Arg138) in the WH1 tails have >10% membrane contact probabilities, and all other residues in this region are below this threshold. Because N-WASP also have a larger number of basic residues in the WH1 tail than WASP, the same contrast between the two proteins described above for the BRs is repeated here. The probability that the WH1 tail is membrane associated is 89% for N-WASP, higher than the 73% for WASP. Likewise, the average number of simultaneously bound WH1 residues is 2.9 for N-WASP, higher than a 2.0 for WASP, and the average number of PIP_2_ molecules bound to the WH1 is 1.4 for N-WASP, also higher than a 1.0 for WASP. These numbers are similar to the counterparts in BR, indicating nearly equal participation of these two regions in membrane association.

The representative snapshots in Figures 3A and 4A show that WASP and N-WASP are tethered to the membrane through both the WH1 tail and BR. The WH1 tails in both proteins have some tendency to form α-helices. In the zoomed images, WASP Arg148 and Arg151 form salt bridges with a PIP_2_ and a PS, respectively, whereas N-WASP Arg141 and Arg143 each form a salt bridge with a PIP_2_.

Based on the above observation that BR residues interact with the membrane largely independently of each other (Figure 2C), we expect independence between WH1 tail and BR in their association with the membrane. To test this idea, we counted the fractions of MD frames for four states: both regions are membrane-bound; only one of the two regions is bound; and neither region is bound. If the regions have membrane association probabilities of *P*_W_ and *P*_B_ on their own, then the independence model predicts the probabilities of the four states as *P*_W_*P*_B_, (1 – *P*_W_)*P*_B_, *P*_W_(1 – *P*_B_), and (1 – *P*_W_)(1 – *P*_B_), respectively. These predictions agree very well with the statistics from the MD simulations (Figure S2), confirming the independence of WH1 tail and BR in membrane association. The best fits yield *P*_W_ = 73% and *P*_B_ = 83% for WASP and *P*_W_ = 89% and *P*_B_ = 92% for N-WASP. These values once again confirm the stronger membrane association of N-WASP relative to WASP, as well as nearly equal contributions of WH1 and BR to membrane association in each protein.

### N-WASP, but not WASP, can simultaneously bind PIP_2_ and soluble Cdc42

We next simulated the membrane association of WASP and N-WASP when they are prebound to soluble Cdc42. Given the independence shown above between WH1 and BR in membrane association, in this set of simulations we left out the WH1 tails. The WASP and N-WASP fragments contained residues Leu195 to Lys288 and Leu157 to Lys253, respectively. They include a portion of the WH1-BR linker and extend to the end of the GBD residues found in the NMR structure of WASP bound with Cdc42 (Protein Data Bank entry 1CEE) [22]. For these simulations, the initial structure of WASP residues Lys230 to Lys288 in complex with Cdc42 (residues 1-179) was taken from 1CEE; the counterpart for N-WASP was built by homology modeling.

A striking difference between the WASP and N-WASP fragments prebound with soluble Cdc42 is that the former does not stably associate with the PIP_2_-containing membrane whereas the latter does. Cdc42-bound WASP only transiently come into contact with the membrane, interrupted by long excursions away from the membrane surface. Not a single BR residue crosses the 10% threshold in membrane contact probability (Figure 2A). In contrast, Cdc42-bound N-WASP is always bound or close to the membrane; the membrane contact probabilities of six BR residues are >10%: Lys189, Lys190, Lys192, Lys194, Lys195, and Lys196 (Figure 2B). This contrast can also be seen by comparing the probability distributions in *n*, the number of BR residues bound to the membrane at a given time (Figure S3A, B). The probability distribution for Cdc42-bound WASP suffers a sharp shift toward *n* = 0, with the total probability for *n* 1 reduced to 20%, from 83% in the absence of Cdc42. Cdc42-bound N-WASP also experiences some loss in the total probability for *n* 1, but the loss is much more modest, down to 62% from 94%. The probability distributions in *m*, the number of BR-bound PIP_2_ molecules, show a similar difference between Cdc42-bound WASP and Cdc42-bound N-WASP (Figure S3C, D).

The reason that Cdc42 prebinding compromises WASP-membrane binding is that the BR is also involved in Cdc42 binding [22], and thus PIP_2_ must compete against Cdc42 for BR binding. In a representative snapshot of the Cdc42-bound WASP simulations (Figure 3B), the BR is away from the membrane. Instead, BR residue Lys232 forms a salt bridge with Cdc42 Glu178 and BR residue Lys235 interacts with two Cdc42 residues, Glu171 via a salt bridge and Asn167 via a hydrogen bond (Figure 3B, zoomed view). There are just too few BR basic residues that are free from Cdc42 interference and thus competent for PIP_2_ binding.

For N-WASP, despite the engagement of some BR basic residues with Cdc42, there are enough BR residues remaining free to make membrane binding stable. A representative snapshot of the Cdc42-bound N-WASP simulations (Figure 4B) illustrates this scenario. BR residue Lys200, which aligns with WASP Lys235 (Figure 1A), form the same interactions with Cdc42 Glu171 and Asn167 as found for WASP (Figure 4B zoomed view, left). In addition, BR residue Lys195 hydrogen bonds with the carbonyl of Gly47. However, the very next residue, Lys196 forms a salt bridge with a PS (Figure 4B zoomed view, right). In the upstream, Lys190 forms a salt bridge with a PIP_2_. In addition, Lys188 and Lys192 stand ready to contact lipids.

### PIP_2_ binding of WASP is restored upon Cdc42 prenylation

Endogenous Cdc42 is prenylated at the C-terminus and inserted into membranes. To model this situation, in our last set of simulations we appended nine residues, P_180_EPKKSRRC_188_, to the C-terminus of soluble Cdc42, added a geranylgeranyl group to the new terminal Cys residue, and inserted the geranylgeranyl group into the hydrophobic layer of the membrane.

Upon Cdc42 prenylation, membrane contact probabilities of WASP recover to the levels without Cdc42 (Figure 2A). The number of BR residues with >10% membrane contact probabilities is now 7; only one residue, Lys235 (see below), is missing from the corresponding list for WASP membrane binding in the absence of Cdc42. The restoration of WASP’s membrane association propensity is corroborated by the probability distributions in the number of membrane-bound BR residues and the number of BR-bound PIP2 molecules (Figure S3A, C), showing near matching without Cdc42 and when WASP is prebound to prenylated Cdc42.

The four basic residues, especially Arg186 and Arg187, in the C-terminal extension are known to contribute to the membrane association of prenylated Cdc42 [35, 36]. In our MD simulations, these residues form extensive salt bridges and hydrogen bonds with PIP_2_ and PS headgroups (Figures 3C and 4C). These interactions, along with the insertion of the geranylgeranyl group into the membrane hydrophobic region, firmly tether Cdc42 to the membrane. Occasionally, some other residues of Cdc42 also come into contact with the membrane, including Lys107, which in the snapshot in Figure 3C forms a salt bridge with a PIP_2_ molecule.

In the presence of prenylated Cdc42, the WASP BR residue Lys235 still forms a salt bridge with Cdc42 Glu171 and a hydrogen bond with Cdc42 Asn167 (Figure 3C zoomed view, right). However, moving upstream, the very next basic residue, Lys232, is already freed from Cdc42 and has a high membrane contact probability (Figure 2A). The membrane-tethered Cdc42 apparently enhances the membrane contact probabilities of the upstream BR residues, making up for the loss at Lys235. One way to achieve this enhancement is presented when basic residues in the Cdc42 C-terminal extension and in the WASP BR bind to the same PIP_2_ molecule. This scenario is seen in the representative snapshot in Figure 3C; in the zoomed view, left, Cdc42 Lys183 and WASP Lys226 bind to the same PIP_2_ molecule. In essence, the Cdc42 C-terminal extension recruits PIP_2_ molecules and raise their local concentration, therefore making it more likely for nearby BR residues to bind to these PIP_2_ molecules. In the same representative snapshot, WASP Arg227 forms a salt bridge with a PS and Lys225 stands ready for alternative or additional membrane contacts. A downstream residue, Arg273, also frequently binds the membrane (Figures 2A and 3C).

In contrast to the dramatic stabilization of the membrane association of the WASP BR, Cdc42 prenylation has little effect on the already strong membrane association of the N-WASP BR found when Cdc42 was in the soluble form (Figure 2B). The probability distributions in the number of membrane-bound BR residues and the number of BR-bound PIP2 molecules also look similar whether Cdc42 is in the soluble or prenylated form (Figure S3B, D).

Representative snapshots of N-WASP prebound with soluble and prenylated Cdc42 in Figure 4B, C illustrate their similarity in membrane association. With either soluble or prenylated Cdc42, a conserved Lys, Lys200, forms the same interactions with Cdc42 Glu171 and Asn167 (Figure 4B zoomed view, left; Figure 4C zoomed view, right) as found for WASP. In N-WASP prebound with prenylated Cdc42, the next upstream basic residue, Arg197, forms a salt bridge with Cdc42 Glu181 (Figure 4C zoomed view, right), instead of binding with the membrane, reminiscent of the hydrogen bonding of Lys195 with the Cdc42 Gly47 carbonyl when Cdc42 was in the soluble form. As found for WASP, the N-WASP BR can cooperate with the C-terminal extension of prenylated Cdc42 in membrane binding, as illustrated by the binding of Cdc42 Lys183 and N-WASP Lys194 to the same PIP_2_ molecule (Figure 4C zoomed view, left). However, this cooperation is not essential for the N-WASP BR, as it has more upstream basic residues for membrane binding. For example, in the same snapshot, Lys190 forms a salt bridge with a PS. A small difference between membrane association in the presence of soluble and prenylated Cdc42 is that, in the latter case, two downstream residues, Lys247 and Arg249 (Figures 2B and 4C) can sometimes contact the membrane.

In short, Cdc42 prenylation has a dramatic effect on WASP-membrane association, making it as strong, or perhaps even stronger, than that in the absence of Cdc42. However, N-WASP behaves similarly in membrane association when it is prebound to either soluble or prenylated Cdc42.

## Discussion

Using MD simulations, we have visualized how WASP and N-WASP bind to both Cdc42 and PIP_2_ to achieve full activation. The simulations recapitulated the puzzling experimental observation of the contrasting effects of Cdc42 prenylation: it is absolutely required for WASP activation but makes no difference in N-WASP activation. In the presence of soluble Cdc42, the WASP BR loses the ability to bind PIP_2_; Cdc42 prenylation recovers this ability. In contrast, the N-WASP BR binds PIP_2_ equally well whether Cdc42 is soluble or prenylated.

The reason of the contrasting effects of Cdc42 prenylation can be attributed to the number of basic residues in BR and how close they are positioned relative to the GBD. GBD-proximal BR residues participate in Cdc42 binding; therefore there is potential for competition between Cdc42 binding and PIP_2_ binding. In the WASP BR, the number of basic residues is lower (7 compared to 10 in the N-WASP BR). In the presence of soluble Cdc42, there are just too few BR basic residues that are free from the interference of Cdc42 binding to sustain stable membrane binding. However, upon Cdc42 prenylation, basic residues in the Cdc42 C-terminal extension and the WASP BR cooperate in binding PIP_2_. Specifically, the basic residues in membrane-tethered C-terminal extension recruits PIP_2_ molecules and prime them for BR binding. In comparison, in the N-WASP BR, the basic residues are not only higher in number but also more spread out from the GBD (Figure 1A). Therefore, there are enough GBD-distal BR basic residues to sustain membrane binding even when N-WASP is prebound to Cdc42 (either soluble or prenylated); some of these residues could cooperate with the C-terminal extension of Cdc42 in PIP_2_ binding when Cdc42 is prenylated.

Given the foregoing understanding, we can speculate distinct mechanisms for the two homologous proteins when prenylated Cdc42 is presented in cellular membranes (Figure 5). Autoinhibited WASP would target its BR toward PIP_2_ molecules enriched around the C-terminal extension of Cdc42 (Figure 5A). Binding of the BR to PIP_2_ molecules would release the C-terminal A motif while binding of the GBD to Cdc42 would release the C-terminal C motif. Because this process requires coordination between PIP_2_ and Cdc42 in a single step, the free energy barrier would be relatively high (Figure 5B). In contrast, the BR of N-WASP would bind to PIP_2_ molecules anywhere in the cellular membrane, resulting in an intermediate where the A motif is released while the C motif is still latched onto the GBD (Figure 5C). Subsequent binding with Cdc42 would free the C motif. In this sequential binding process, the separate free energy barriers are relatively low (Figure 5D).

**Figure 5.**
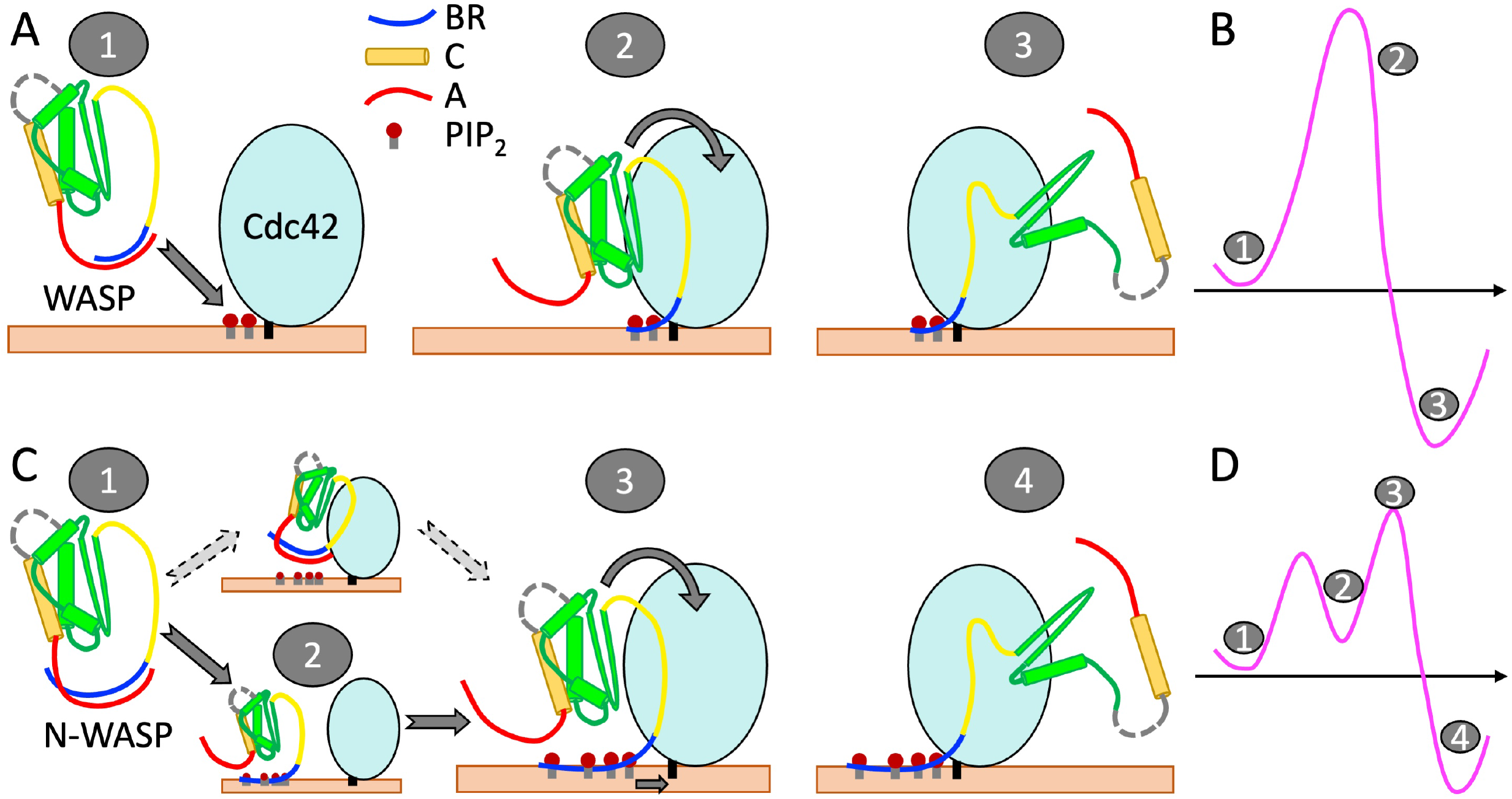
Distinct mechanisms of WASP and N-WASP activation. (A) Activation of autoinhibited WASP starts with targeting its A-bound BR to PIP_2_ molecules around the C-terminal extension of prenylated Cdc42 (state “1”). In a single concerted step, PIP_2_ molecules peel the BR away from the A motif (state “2”) and Cdc42 wrestles the GBD away from the C motif, leading to the activate state (“3”). (B) Free energy diagram of the WASP activation pathway. Note that state 2, in which PIP_2_ has replaced the A motif as the interaction partner of the BR, is just beyond the free energy barrier. (C) Activation of autoinhibited N-WASP (state “1”) starts with PIP_2_ displacing the A motif as the interaction partner of the BR, producing an intermediate state (“2”). (An alternative pathway, where N-WASP first targets Cdc42, should have a much higher free energy barrier and thus is much less likely.) Subsequently, PIP_2_-bound N-WASP engages with prenylated Cdc42 (state “3”). The complete binding of Cdc42 with the GBD and the proximal portion of the BR completes the activation (state “4”). Note that N-WASP has a longer BR than WASP and a correspondingly longer A motif. (D) Free energy diagram of the N-WASP activation pathway.

The simulation results for the simultaneous interactions of WASP and N-WASP with PIP_2_ and Cdc42 may prove insightful for the activation of other IDPs that require multiple regulators. One advantage for having multiple regulators is that they produce sharp, or switch-like response. The sharpness increases with the degree of cooperativity between the regulators. Thus we expect that, other things being equal, the activation of WASP would be more switch-like than that of N-WASP. However, the sharper response comes at a cost: to overcome the corresponding higher free-energy barrier would require higher levels of Cdc42 or PIP_2_. This cost could be a contributing factor to why the expression of WASP is restricted to hematopoietic cells whereas the expression of N-WASP is ubiquitous.

Comparing the sequences of WASP and N-WASP, the former has a ∼30-residue insertion in the linker between WH1 and BR whereas the latter has an additional V toward the C-terminus. Both of these insertions have direct functional consequences. The additional V motif in N-WASP increases the activity for stimulating actin polymerization [16], whereas the 30-residue insertion in WASP is responsible for the chemotaxis of T-cells toward a chemokine [11]. Here we show that even subtle sequence differences in a shared region, the BR, cause drastic disparity in the activation of the two homologous proteins, specifically in the effects of soluble Cdc42 and the effects of Cdc42 prenylation.

As noted above, one aspect of the differences in the BR sequences concerns the spacing of the basic residues from the GBD. Veltman and Insall [12] observed that the shift of basic residues toward the GBD is an evolutionary trend. This trend is evident when the 44 amino acids preceding the CRIB motifs of an ancestral (*<u>Dictyostelium discoideum</u>*, or Dd) WASP, invertebrate N-WASPs, vertebrate N-WASPs, and human WASP are lined up (Figure S4A). We can quantify this trend by using ReSMAP [30] to predict the membrane binding profiles of these sequences (Figure S4B). By fitting each profile to a Gaussian distribution, we identify the peak as the BR centroid and calculate its spacing from the start of the CRIB motif. The spacing is 20 for Dd WASP, then exhibits a sharp decline to 12 at the emergence of invertebrates, and finally taper off to 9 for human WASP (Figure S4C). It will be interesting to use this wide set of WASP orthologs to validate the expected effects, based on the present study, of the BR centroid-CRIB spacing on WASP activity. For example, the wide spacing from GBD is expected to allow Dd WASP BR to bind PIP_2_ largely independent of GBD-Cdc42 interactions.

Functional studies have indeed shown that this BR is crucial for Dd WASP to localize at sites rich in PIP_2_ (and PIP_3_) and stimulate actin polymerization during chemotaxis [37].

## Computational Methods

### System preparation

To prepare for membrane association simulations, WASP and N-WASP fragments consisting of residues 133-255 and 116-220, respectively, were assigned initial conformations using TraDES [38]. These fragments were first simulated in water (with 100 mM NaCl) for 500 ns in quadruplicate with different random number seeds on GPUs using pmemd.cuda [39] in AMBER18 [40]. Following previous studies [41, 42], the force fields were ff14SB [43] for proteins and TIP4P-D [44] for water. Four snapshots each for WASP and N-WASP were taken from these simulations to be used as starting structures for membrane association simulations without Cdc42.

For membrane association simulations with Cdc42, the WASP and N-WASP fragments contained residues 195-288 and 157-253, respectively. The initial structure of WASP residues 195-229 was taken from a snapshot in a simulation without Cdc42; the remaining WASP residues and Cdc42 residues 1-179 were taken from 1CEE [22]. The initial structure of N-WASP prebound to Cdc42 was generated in a similar way, but with homology modeling using 1CEE as the template. Prenylated Cdc42 was generated by appending P_180_EPKKSRRC_188_ to the C-terminus of Cdc42 from 1CEE, modifying the Cys188 sidechain with a geranylgeranyl group, and adding a methyl group to the backbone terminal carboxyl group. The proteins were placed atop a membrane and solvated using CHARMM-GUI [45]. The upper leaflet contained 400 lipids, with the following composition: PIP_2_, 10%; POPS, 40%; POPC, 20%; POPE, 20%; cholesterol, 10%. In the lower leaflet, PIP_2_ and POPS were absent and replaced by POPC (at a slightly smaller number, because the surface area of POPC is ∼10% larger than that of POPS). The geranylgeranyl group of prenylated Cdc42 was inserted into the upper leaflet. Na^+^ and Cl^−^ ions were added to neutralize each system and make up 150 mM salt. The lipid force field was Lipid17 [46]. Each system contained ∼430,000 atoms.

### Molecular dynamics simulations

Each system with the initial structure prepared above was first equilibrated in NAMD 2.12 [47] following a six-step protocol [45], in which, after 10,000 steps of conjugate-gradient minimization, constraints on protein and lipid atoms were gradually released. The timestep was 1 fs in the first three steps (two steps of 25 ps each at constant NVT and 1 step of 25 ps at constant NPT), and increased to 2 fs for the remaining steps of 100 ps each at constant NPT. All bonds involving hydrogens were restrained using the SHAKE algorithm [48]. The nonbonded cutoff was 12 Å (with force switching started at 10 Å for van der Waals interactions). Long-range electrostatic interactions were treated by the particle mesh Ewald method [49]. Temperature was maintained at 300 K using the Langevin thermostat [50] with a damping constant of 1.0 ps^-1^. Pressure was maintained at 1 atm by the Langevin piston [51] with an oscillation period of 50 ps and decay time of 25 ps.

The simulations were then continued in quadruplicate on GPUs using pmemd.cuda [39]. Pressure was now regulated by the Berendsen barostat [52]. Semi-isotropic scaling was applied in the x-y direction to maintain the overall shape of the system. In total, WASP and N-WASP simulations without Cdc42 were run in 16 replicates, each lasting 1360 ns; WASP simulations with soluble and prenylated Cdc42 were run in 4 copies, each lasting 1000 ns; and N-WASP simulations with soluble and prenylated Cdc42 were run in 4 copies, each lasting 700 ns. The first 360 ns of the simulations without Cdc42 and the first 100 ns of the simulations with Cdc42 were discarded. The remaining portions were saved at 20 ps intervals for analysis.

## Data analysis

Contacts between protein and lipid heavy atoms were calculated in cpptraj [53] and further analyzed using mdtraj [54]. A contact was defined between two heavy atoms within 3.5 Å.

## Acknowledgments

This work was supported by National Institutes of Health Grant R35 GM118091.

## Author contributions

SD: investigation, methodology, writing - original draft. H-XZ: conceptualization, funding acquisition, supervision, writing - original draft, writing - review & editing.

