## Supplementary Figures for "Why Does Synergistic Activation of WASP, but Not N-WASP, by Cdc42 and PIP_2_ Require Cdc42 Prenylation?"

*Supplementary Information*

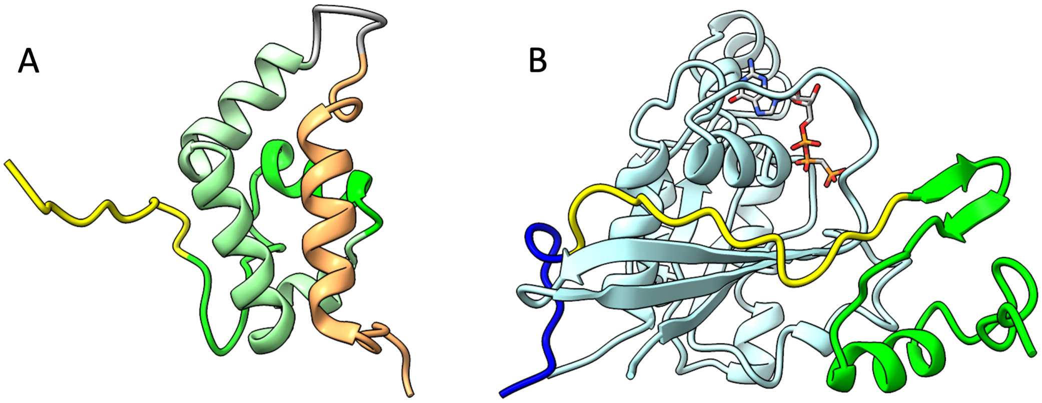


Figure S1. NMR structures of the WASP GBD in autoinhibited and activated states. (A) GBD bound with the C motif, from Protein Data Bank entry 1EJ5. CRIB and C motif are in yellow and orange; the GBD portion that binds to Cdc42 is in green, whereas the remaining portions are in two successively dimmer shades of green. (B) GBD bound to Cdc42. In addition to the colors for the regions present in (A), BR is in blue and Cdc42 is cyan; a GTP is shown as stick.

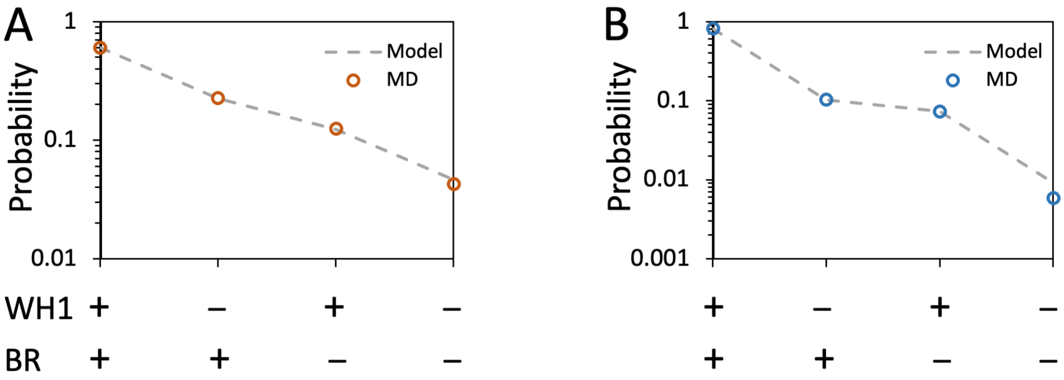


Figure S2. Comparison of WH1 and BR membrane association probabilities with the predictions assuming independence between the two regions in membrane association. (A) WASP. (B) N-WASP. The statistics obtained from MD simulations are shown as circles and model predictions are shown as dashed lines, for four possible states (bound and unbound denoted by + and – signs).

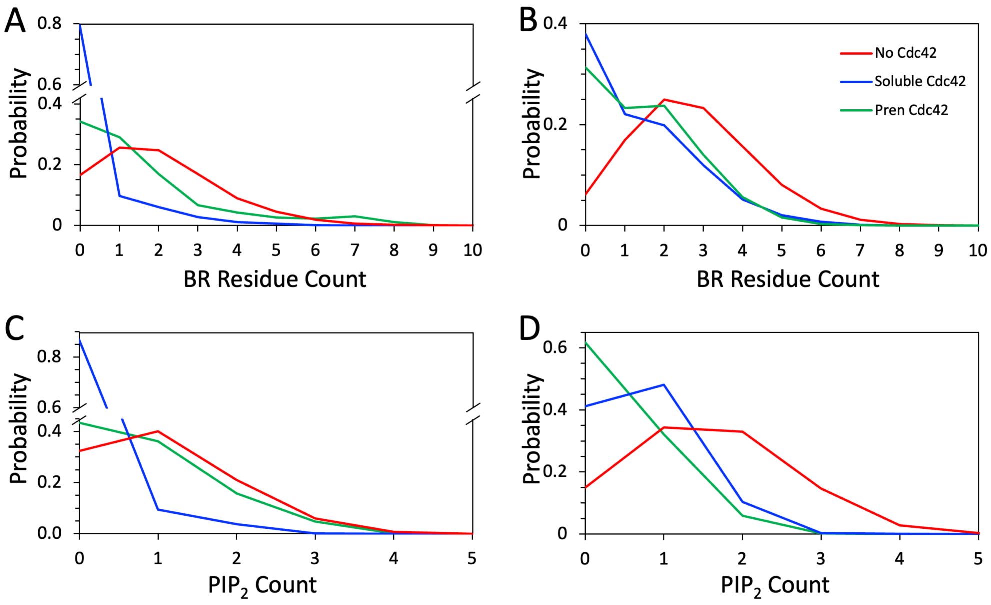


Figure S3. Probability distributions for *n* BR residues simultaneously bound to membranes and for *m* $PIP_2$ lipids simultaneously bound to the BR. (A) Distributions for *n* (BR residue count) in WASP simulations. (B) Corresponding plots for N-WASP. (C) Distributions for *m* ($PIP_2$ count) in WASP simulations. (D) Corresponding plots for N-WASP. Results from simulations without Cdc42, with soluble Cdc42, and with prenylated Cdc42 are in red, blue, and green, respectively.

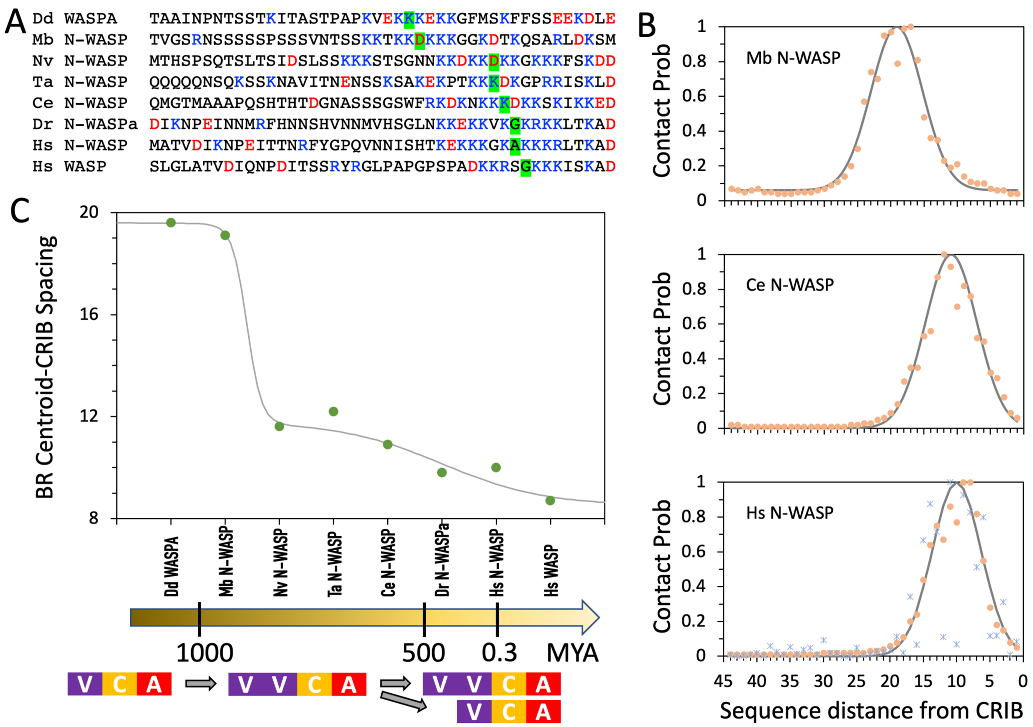


Figure S4. Spacing between BR centroid and CRIB in WASP orthologs. (A) Sequence comparison for 44 amino acids immediately before CRIB in eight WASP orthologs. The amino acid closest to the BR centroid identified in (B) is shaded in green. Abbreviations (and UniProt entry names) are: Dd, *Dictyostelium discoideum* (Q9GSG9); Mb, *Monosiga brevicollis* (A9V6R1); Nv, *Nematostella vectensis* (A7RXK9); Ta, *Trichoplax adhaerens* (B3RTH0); Ce, *Caenorhabditis elegans* (Q8MQE6); Dr, *Danio rerio* (F1QNY1); Hs, *Homo sapiens* (O00401 for N-WASP and P42768 for WASP). (B) Membrane association propensities predicted by ReSMAP [30] from selected sequences in (A), shown as orange circles; Gaussian fits are shown as solid curves. The peak position of each Gaussian is identified as the BR centroid. For Hs N-WASP, the membrane contact probabilities from the MD simulations, scaled to a maximum of 1, are shown as blue asterisks. (C) The progression of the BR centroid-CRIB spacing over evolutionary time (million years ago, or MYA). The curve is only for guiding the eye. Invertebrates emerged shortly after 1000 MYA; vertebrates emerged ~500 MYA; humans emerged 0.3 MYA. The classification into N-WASP is based on the presence of an extra V motif. According to Veltman and Insall [12], metazoan N-WASPs descended from ancestral WASPs, whereas vertebrate WASPs branched off from this lineage.
